# Environmental filtering, dispersal limitation, and competition control the distribution of acidophilic iron oxidizers

**DOI:** 10.64898/2026.04.22.720147

**Authors:** Christen L. Grettenberger, Caden Williams, Trinity L. Hamilton

**Affiliations:** Department of Earth and Planetary Sciences, University of California, Davis; Department of Environmental Toxicology, University of California, Davis; Department of Plant and Microbial Biology and The BioTechnology Institute University of Minnesota

## Abstract

Acid mine drainage is a global pollution problem characterized by low pH and high concentrations of metals. Active remediation is often cost-prohibitive, but Fe(II) oxidizing microbes may be used for passive bioremediation. To leverage these species, we must understand the factors that control their distribution. Here, we examine the environmental and ecological factors that control these species with the aim of determining if microbial seeding is a viable remediation strategy. Although stochastic processes appear to control the distribution of majority of taxa inhabiting AMD ecosystems, the distribution of Fe(II) oxidizers is driven by environmental filtering and competition. The abundance of all the major Fe(II) oxidizing genera have significant relationships with pH, with pH explaining 10 – 38% of the variation in their abundance. The genera appear to have pH preferences with *Acidithiobacillus* and *Leptospirillum* preferring environments below pH 3, *Gallionella, Sideroxydans*, and *Ferritrophicum* preferring environments above pH 3.5, and *Ferrovum* preferring intermediate pH environments. Once the effect of pH is removed, genera that share pH preferences are negatively correlated, indicating that they are likely competing for the Fe(II) oxidizing niche in their preferred environments. Communities are also shaped by dispersal limitation, which suggests that microbial seeding is possible in these environments. Future seeding attempts should consider species interactions and ecology more generally to inform their efforts.

## Introduction

Acid mine drainage (AMD) is a major environmental problem that pollutes watersheds across the globe [1–5]. It is created when iron mining activities expose iron-sulfur mineral bearing rocks generating runoff with low pH and a high concentration of sulfate and metals, including Fe, Cu, and As. Bioremediation is often used in AMD ecosystems because active remediation is extremely expensive [6]. Unfortunately, in low-pH systems, many traditional methods of bioremediation fail. For example, constructed wetlands do not function below pH 4, and limestone-lined channels become armored by iron oxides and therefore have short useful lifetimes [6]. One promising methodology leverages the naturally occurring microbial communities. In these systems, the environment is engineered for microbial communities to oxidize Fe(II), causing the formation of iron oxides. These iron oxides sorb other metals, effectively removing them from the drainage. Downstream, acidity can be removed using limestone-lined channels without the risk of armoring [6]. This system hinges on Fe(II) oxidizing microbial communities effectively and rapidly removing Fe(II). Iron oxidizing bacteria oxidize Fe(II) at different rates and under different environmental conditions [7]. Thus, it may be beneficial to seed the most effective Fe(II) oxidizers to facilitate remediation [8]. However, choosing the proper taxa for use in bioremediation requires not only that those species rapidly oxidize Fe(II), but also that they can successfully colonize and persist in the environment.

Seeding an environment with a microbial community to alter how that environment functions is not a new idea. Microbiome transplants have been used to cure deadly *Clostridium difficile* infections in humans [9– 11] and in attempts to increase the degradation of petroleum products [12], the productivity of crop fields [13], and nitrogen retention in soil mesocosms [14]. In the same vein, seeding AMD ecosystems with the species that rapidly oxidize Fe(II) is a promising solution to improve the rate at which iron and other metals are removed from the system. However, seeding a community can have mixed results. Seeding hydrocarbon-degrading bacteria into petroleum-contaminated environments increased petroleum degradation in some cases but had no effect in others [12] and seeding compost to speed the degradation process altered the community composition but did not influence the rate of organic matter degradation [15]. Part of the variable success in seeding likely results from the species ability to establish themselves in the new environment. Although environmentally based parameters, for example the pH, concentration of oxygen, or nutrient availability, are likely easy to measure and account for, seeding must also account for interspecies interactions. These interactions are difficult to see, much less quantify. Therefore, many of these interactions occur within a black box, making it difficult to select the appropriate microbial community.

The efficacy of seeding efforts requires accounting for the dominant factors that influence microbial community assembly and composition. Community composition can be determined by deterministic processes including environmental filtering and species interactions like competition and cross feeding. Stochastic processes like population drift can also impact community composition as can dispersal limitation. If the processes controlling community assembly in AMD ecosystems are primarily deterministic, then communities should assemble to a single stable community [16, 17] and seeding efforts are unlikely to be effective. If dispersal limitation is a key factor, then communities assemble semi-stochastically [18], and seeding the community may allow the target taxon to establish itself in the community and contribute to metal removal. If the community composition is driven by the initial abundance of species and stochastic processes, then seeding efforts can help to drive the community toward the desired composition.

Microbial community composition in AMD sites appears to co-vary with pH and the concentration of Fe(II) [5, 19, 20]. However, more recent data suggest that these taxa inhabit a wider range of environments [21]. For example, the genus *Ferrovum* was thought to inhabit a niche around pH 2.4 to 3 [5, 19, 20] but it is the predominant taxon in environments with a pH as low as 2.1 [3, 21, 22]. The limitations of these initial niche models may result from those studies examining microbial communities within a single region – the Appalachian Coal Belt [19, 20] and China [5], respectively. Thus, to understand the ecological and environmental factors that control microbial community composition and inform potential seeding efforts, we must examine microbial communities from AMD sites globally.

Here, we examine the microbial communities from 315 AMD environments from across the globe to determine what factors control the distribution and diversity of bacterial species in AMD environments and whether a shared or core set of taxa are found in AMD environments. We use neutral population models to determine if species distributions are controlled by stochastic or deterministic processes and use species co-occurrences to determine how ecological interactions may influence the distribution of Fe(II) oxidizing taxa.

## Methods

We searched the NCBI Sequence Read Archive for AMD samples using the key term “acid* mine” where “*” is a wildcard. It is not possible to compare communities sequenced with different primer pairs, because each primer has a unique bias against certain groups. Therefore, for each bioproject, we identified the primer set used based on the associated publication or information available in NCBI. Only the primer set 515f and 806r had sufficient samples to compare. However, there are multiple versions of the 515f and 806r primer set and it was not always possible to determine if the 515f and 806r primers used were the original or the modified versions. These versions differ in their coverage of the archaea and SAR11 clade [23, 24] SAR11 is a marine taxon and therefore does not comprise a large portion of our community and we removed archaeal sequences (see below). Therefore, we use all samples sequenced using the 515f and 806r primer set. After retrieving the NCBI results, we realized that there were AMD studies using the 515f and 806r primer pair that had not been identified in our search. Therefore, we searched Google Scholar using the search terms “acid mine drainage” AND “515f” AND “806r.” We examined each result to determine if they included 16S rRNA amplicon datasets sequenced with Illumina sequencing and using the 515f and 806r primer set. If so, the samples were added to the dataset.

We gathered environmental and geochemical data from each of the papers associated with the data gathered above. We removed samples that were from bioreactors, active treatment systems, or cultivated samples, those above pH 5, and those for which we could not retrieve geochemical data. We also removed five samples from one bioproject [25] because these samples generated contigs that were not the correct length. The bioprojects used for this work are listed in Table 1.

**Table 1.**
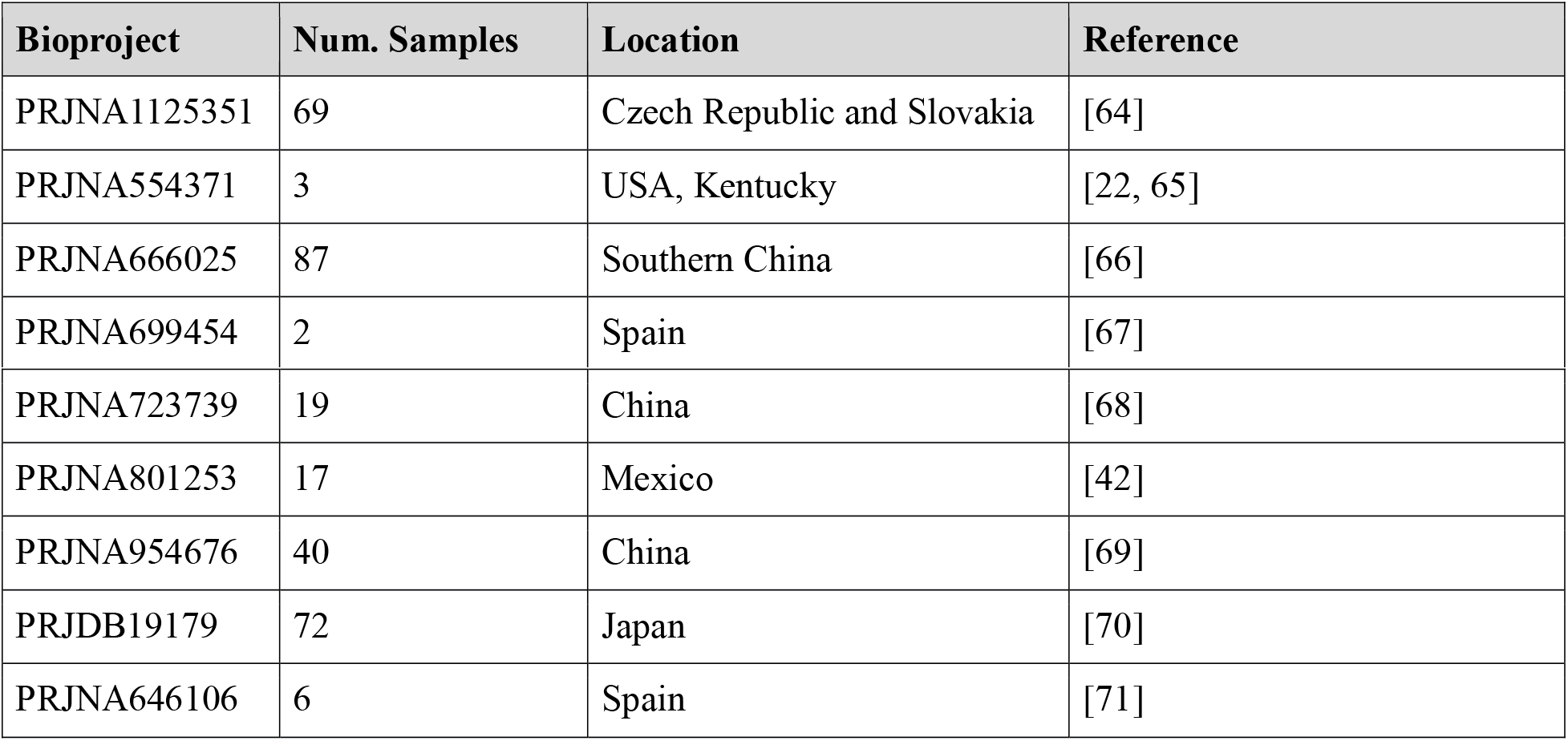
Bioprojects used in this analysis, the number of samples used per project, the sampling location, and the reference.

| Bioproject | Num. Samples | Location | Reference |
| --- | --- | --- | --- |
| PRJNA1125351 | 69 | Czech Republic and Slovakia | [64] |
| PRJNA554371 | 3 | USA, Kentucky | [22, 65] |
| PRJNA666025 | 87 | Southern China | [66] |
| PRJNA699454 | 2 | Spain | [67] |
| PRJNA723739 | 19 | China | [68] |
| PRJNA801253 | 17 | Mexico | [42] |
| PRJNA954676 | 40 | China | [69] |
| PRJDB19179 | 72 | Japan | [70] |
| PRJNA646106 | 6 | Spain | [71] |

All data sets were analyzed in *dada2* (v. 1.36.0) [26]. Because the error rate estimation calculated by *dada2* (in the *learnerrors* and *dada* steps) requires that the samples analyzed together were sequenced in the same run, samples from each bioproject were assumed to have been sequenced in the same sequencing run and were processed together [26]. Samples from different bioprojects were analyzed separately through the chimera removal step (see below). For example, the 69 samples from bioproject PRJNA1125351were sequenced together and separately from the three samples from bioproject PRJNA554371. For each bioproject, sequences were checked to determine if they contained primer sequences using degenerate versions of 515f (GTGNCAGCMGCCGCGGTAA) and 806r (GGACTACNVGGGTWTCTAAT) as the search term. These degenerate sequences were designed by combining all known modified versions of the 515f and 806r primers. If the primers were present, then they were removed using trimleft in the *filterandtrim* command. Sequences were also trimmed based on their quality profile and sequences with N’s, those with more than 2 expected errors, and those that were identified as phiX were removed, also with the *filterandtrim* command. Sequencing error rates were estimated using the *learnerrors* command and were applied using the dada command. Forward and reverse reads were merged using the *mergePairs* command. Samples for which most reads did not merge were removed. Sequences that were shorter than 245 bp or longer than 260 bp were removed. Putatively chimeric sequences were identified and removed using the *removeBimeraDenovo* command. Then, samples from all bioprojects were combined into a single sequence table using the *mergeSequenceTable* command. Sequences that were identical except for length (i.e. differed due to trimming) were collapsed using the *collapseNoMismatch* command. Sequences were classified using the *assignTaxonomy* command using the Silva non-redundant database (v. 138.2) [27]. A *phyloseq* object was created using the sequence table, the classifications, and the environmental and geochemical data using the R package *phyloseq* (v. 1.52.0) [28]. Sequences identified as Chloroplasts or Mitochondria and those not classified as bacteria were removed using the *subset_taxa* command in *phyloseq*. Huang’s Coverage was calculated using the *coverage* command in the R package *entropart* (v 1.6.16) [29].

The core or shared microbial community was defined as genera or ASVs that are present in >75% of samples. This analysis was done at both the ASV and genus level. We chose to perform a genus level core community because core metabolisms are often shared at the genus level and because individual species appear be restricted to individual sites or regions [30]. Genera were grouped using the *glom_taxa* command in *phyloseq*. ASVs and genera that were present in >75% of the samples were considered the core community.

Diversity metrics including richness and Shannon and Simpson diversity were calculated for the non-grouped (i.e. ASV) and the genus grouped datasets using the *estimate_richness* command in *phyloseq*. We calculated the correlation between pH and richness and diversity using the *lm* command in RStudio (v 2025.05.0+496) [31, 32].

We generated a neutral model for community assembly using the *neutral*.*fit* command in the R package *MicEco* (v. 0.9.19) [33] which uses the model developed by Sloan and others [34] and implemented by Burns and others [35]. Here, we report on the goodness of fit (R^2^) and immigration rate (m).

Because Fe(II) oxidizing species are of particular interest in AMD ecosystems, we sought to understand the distribution of well-known Fe(II) oxidizing genera *Leptospirillum, Acidithiobacillus, Ferrovum, Gallionella, Ferrotrophicum*, and *Sideroxydans*. However, this approach does have limitations including a lack of comprehensive understanding of the metabolic potential of all species within these genera. For example, some *Acidithiobacillus* oxidize sulfur rather than iron (e.g., *A. thiooxidans*) or can oxidize or reduce other metals [45, 46]. And 16S rRNA sequences often do not classify to the species level. Each of these genera is known to have Fe(II) oxidizing species that are common in AMD or neutral iron rich environments [5, 20, 36–44]. Because our NCBI Sequence Read Archive search targeted acid mine drainage data sets, we assume that the recovered taxa could be Fe(II) oxidizing taxa.

We used a generalized additive model (GAM) to model the relationship between the abundance of each genus of interest. Before running the model, counts were transformed into relative abundance. Then we computed the centered log ratio transform using the *clr* function in the R package *compositions* v2.0-8 [47]. Before transformation a pseudocount of 1E -6 was added to avoid zeros. The GAM was calculated using the *gam* function in the R package *mgcv* v1.9-3 [48] using the restricted maximum likelihood approach. k was set to 5 to avoid overfitting the data. The basis argument was set to cubic regression (cs). P-values were adjusted for multiple comparisons using the *p*.*adjust* function in R [32] and using a Bonferroni correction. We determined the optimal pH for each genus of interest by solving the best fit GAM model at 500 pH values and picking the value at which the abundance was greatest. We removed samples that had the lowest and highest 2.5% of pH values when predicting the optimal pH to avoid sparse data at high and low pH values from erroneously influencing the splining at the edges.

We extracted the residuals from the GAM and used a graphical lasso to test for covariance between genera after removing the influence of pH. The graphical lasso [49] was implemented using the *huge* command in the *huge* package v1.3.5 [50] using the glasso method and nlambda of 20. The best model was selected using EBIC. We repeated the GAM modeling for ASVs to determine if individual ASVs of the Fe(II) oxidizing genera had significant relationships with pH. We also used a graphical lasso as described above to determine if the abundance of individual ASVs within a genus were positively or negatively correlated. ASVs that were present in less than 3 samples were removed when constructing the ASV level graphical lasso. Because the genus *Acidithiobacillus* contains species that oxidize sulfur in addition to or instead of iron, we grouped ASVs by species and repeated the GAM modeling for each species.

## Results

315 samples from ten bioprojects met our criteria for inclusion. The only geochemical parameter reported for all samples was pH. After removing non-bacterial sequences, these samples contained 24,197 ASVs with 3 – 3,456 ASVs per sample. The least rich samples (those with fewer than 20 ASVs) were from biostalactites in mines where the pH was 3.0 or below. The richest samples (those with >1,000 ASVs) were from AMD environments with pH 3.95 and above in Japan and China. The median number of ASVs was 149. Huang’s coverage was >99.9% for all samples.

The community included 73 phyla, and each sample contained members of 3 - 46 phyla (median 21). The most abundant phylum was the Pseudomonadota (formerly Proteobacteria) which made up 1.5 – 99.8% of the microbial communities (median 73%) within the Pseudomonadota, the most abundant families were the Gallionaceae, Acidithiobacillaceae, Ferrovaceae, Acidiferrobacteraceae, Acetobacteraceae, Rhodanobacteraceae, Sulfuricellaceae, and Legionellaceae all of which made up > 40% of the microbial community in at least one sample. Other abundant phyla included the Actinomycetota (median 4.4%), Acidobacteriota (median 3.2%), Sva0485 (median 2.6%), Planctomycetota (median 1.6%), and Bacillota (median 0.7%). The Nitrospirota made up 0 – 72% of the microbial community (median 0.7%). Within this phylum, the Leptospirillaceae was the most abundant family and made up >40% of the microbial community in at least one sample.

### Core Microbial Community

Six genera were present in 75% or more of the communities including the gammaproteobacterial genus *Ferrovum*, an unnamed genus within the Acetobacteraceae, the Acidobacteriales genus *Leptospirillum* and two unclassified Actinomycetota genera within the family Acidimicrobiaceae and Acidimicrobiaceae subgroup 1 and an unclassified member of the Bacteria. No ASVs were present in more than 34% of communities and only 46 were present in more than 20% or more of the communities.

### Community Modeling

Most taxa (84.5%) appeared as frequently as predicted by the neutral model, a smaller fraction appeared more frequently than predicted (15.4%) and few appeared less frequently than predicted (0.14%) (Table 2, Figure 1A). The m was 5.10^-5^. The distribution of the predominant Fe(II) oxidizers differed from that of the community as a whole. All Fe(II) oxidizing genera appeared less frequently than predicted based on the model. Individual ASVs within the genera were less likely to fit the neutral model than the community as a whole (Table 2, Figure 1B). There was a significant relationship between pH and observed richness (number of ASVs) (p < 0.01; R^2^ = 0.09) and with Shannon diversity (p < 0.01, R^2^ = 0.03). All Fe(II) oxidizing genera had significant relationships with pH (Table 3, Figure 2A-F). The pH optima for each genus were between 2.18 and 4.5 (Table 3, Figure 2) and the pH optimum for *A. ferrooxidans* was approximately the same as for the genus *Acidithiobaciullus* (2.19 and 2.18 respectively). All species within the *Acidithiobacillus* except *A. ferrovorans* had significant relationships with pH (Supplemental Table 1). However, only *A. ferrooxidans* was abundant in the sample data (average abundance >1%) (Supplemental Table 1).

**Table 2.**
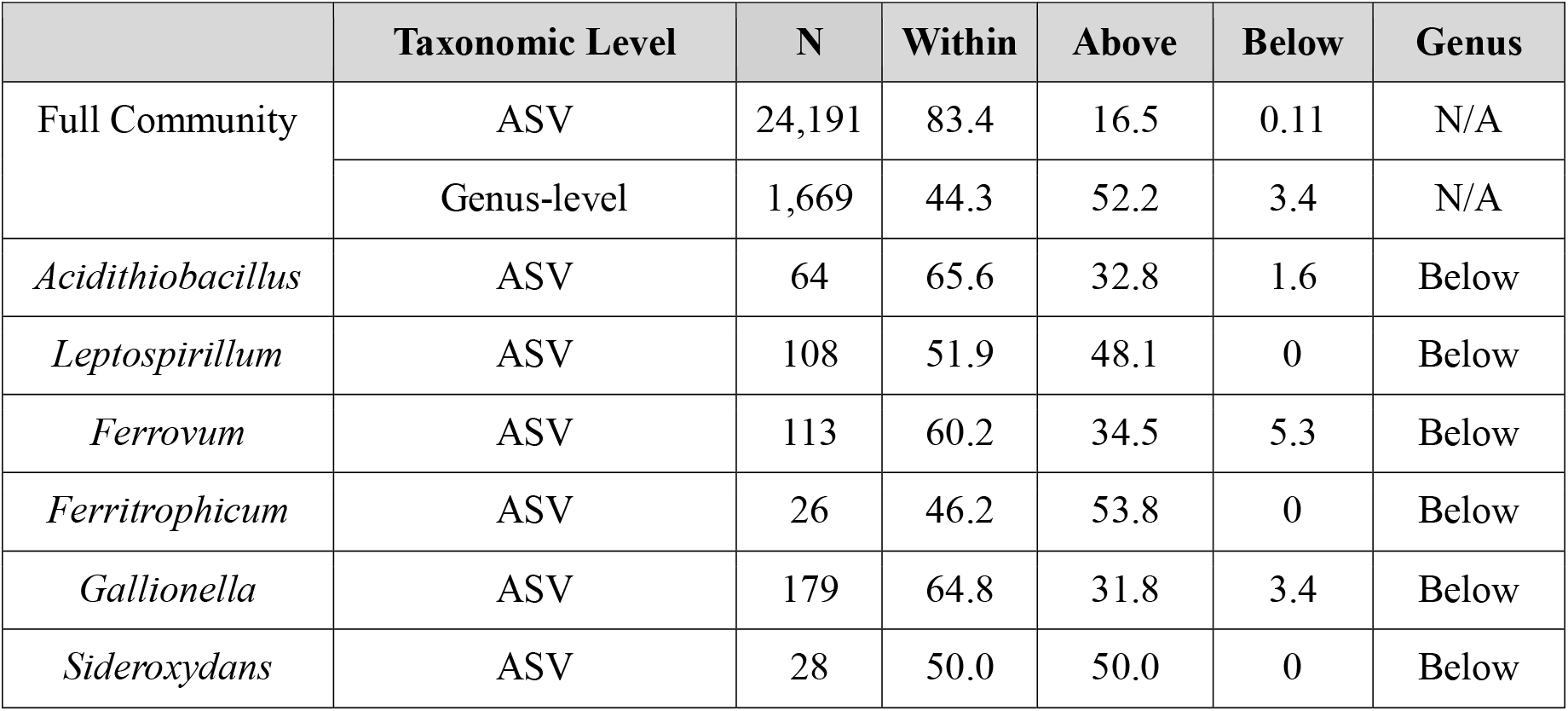
Percentage of ASVs with frequencies that are within, above, and below the frequency predicted by the neutral model. For one analysis, genera, rather than ASVs were used for the analysis. This is indicated in the taxonomic level column. The number of taxa used for the analysis is indicated in the column labeled N. For each genus, the analysis was performed the ASV level (which is reported in the within, above, and below columns). Whether the genus was above, below, or within the frequency predicted in the genus-level analysis is indicated in the column labeled Genus.

**Table 3.**
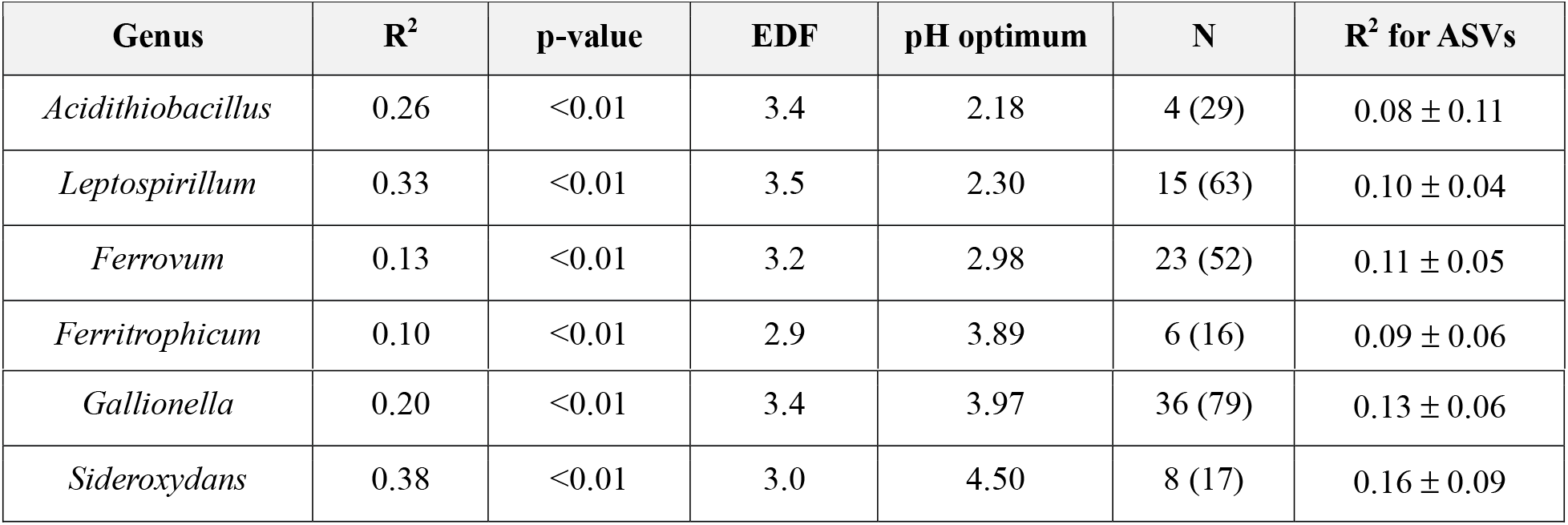
Percentage of variance in the abundance (R^2^) of a genus that is explained by pH using a generalized additive model, the significance of the relationship (p-value) and the estimated degrees of freedom (EDF) for the chosen model. The number of ASVs with a significant relationship with pH (N) with the total number of ASVs in the genus that were modeled in parentheses. The range of R^2^ values for individual ASVs with significant relationships with pH is indicated in the final column. For ASV level analyses, only taxa that were present in 3 or more samples were included.

**Figure 1.**
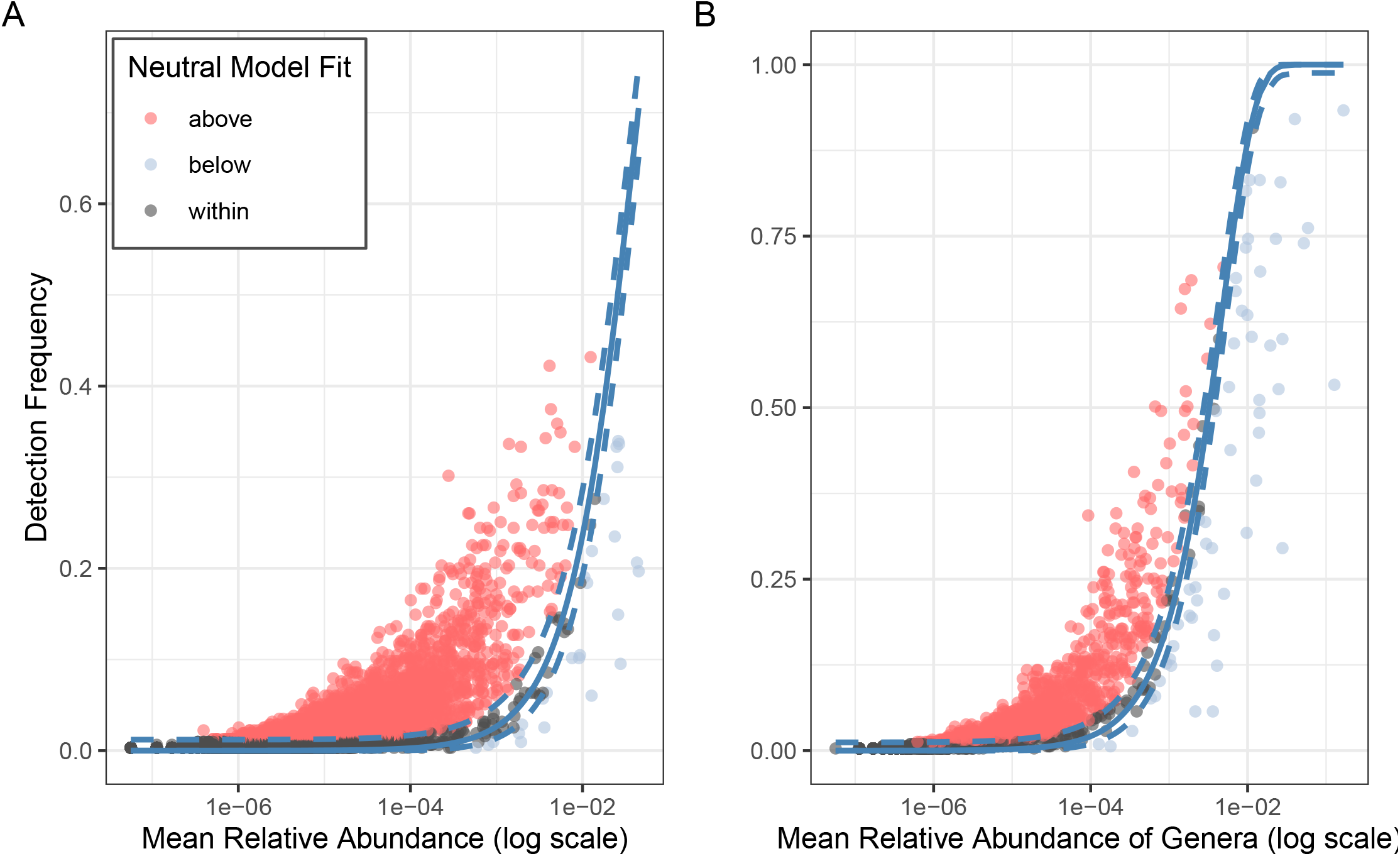
Neutral community model for (A) AMD genera and (B) AMD ASVs. Those that appear more frequently than predicted by the community model are indicated by red, those that appear less frequently by blue, and those that appear as predicted are grey. The neutral model is indicated by the grey line and upper and lower confidence intervals by the dashed line.

**Figure 2.**
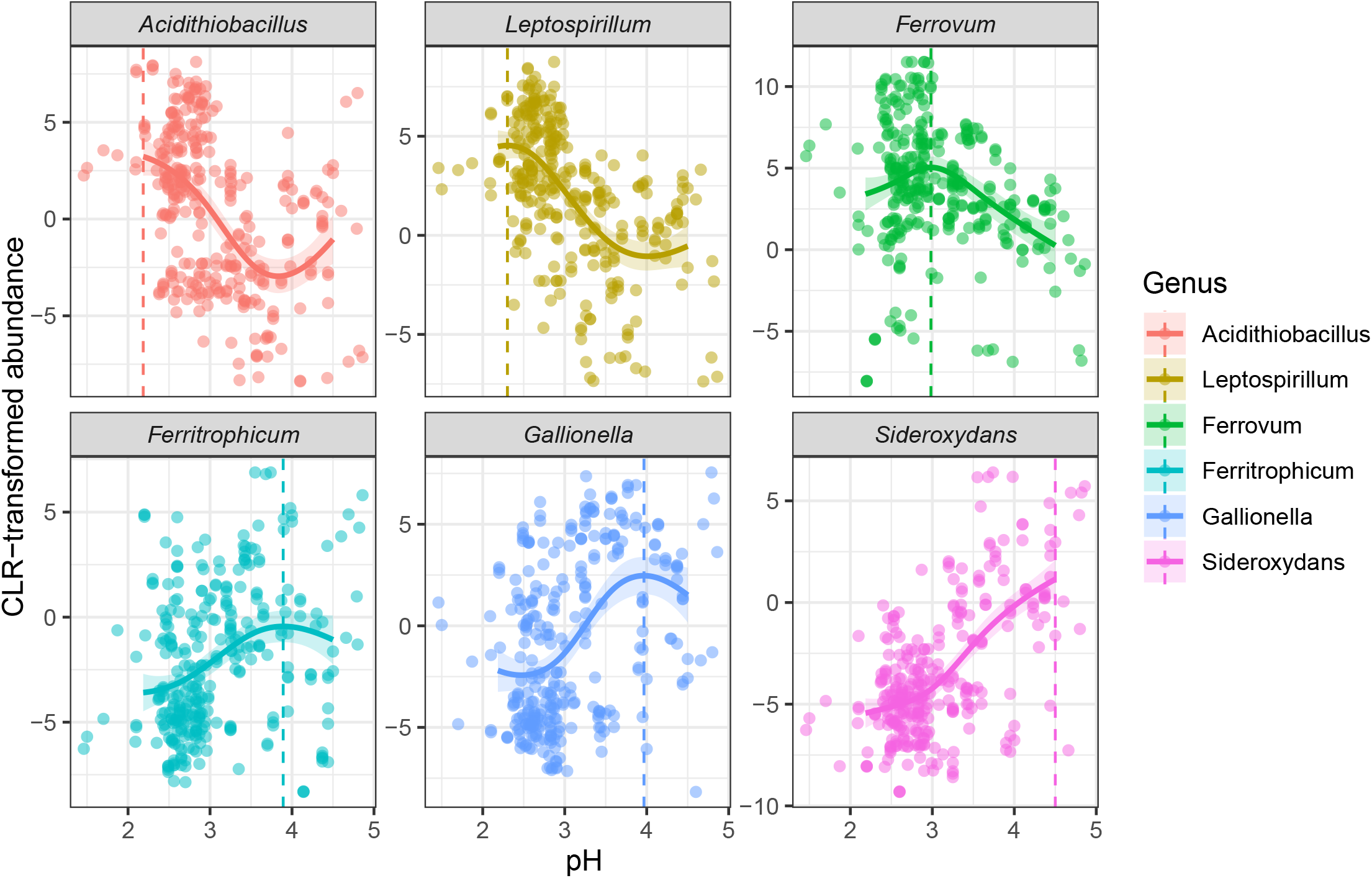
GAM model and CLR adjusted abundances for AMD genera. CLR-transformed abundances are indicated by points, and the GAM model is plotted with a line. The pH optimum is indicated by a dashed line.

After removing the effect of pH, eight genus pairs had non-zero correlation coefficients. *Acidithiobacillus* was negatively correlated with *Leptospirillum*, and *Ferritrophicum*. Similarly, *Gallionella*, and *Sideroxydans* were negatively correlated with one another. *Acidithiobacillus* was positively correlated with *Ferrovum* and *Sideroxydans*. Similarly, Leptospirillum was positively correlated with *Sideroxydans*. Finally, *Ferrovum* was positively correlated with *Ferritrophicum* (Table 4). Species within the same genus primarily had non-significant partial correlation coefficients (Table 5). Of the taxa pairs with non-zero partial correlation coefficients, >90% were negative.

**Table 4.** Precision matrix for the major Fe(II) oxidizing genera. The partial correlation coefficient 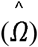 represents the correlation of the two taxa after the effect of all other taxa are removed. Because this analysis was performed on the residuals of the GAM, this represents the partial correlation after all other taxa and the effect of pH have been removed. Taxa with negative correlations are highlighted in orange. Those with positive correlations are highlighted in green. For taxa with non-zero correlation coefficients, the difference in their optimum pH is indicated in parentheses. Graphical lasso does not provide a p-value or an R^2^, thus only the partial correlation coefficient 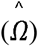 is reported here.

|  | pH | *Acidithiobacillus* | *Leptospirillum* | *Ferrovum* | *Ferritrophicum* | *Gallionella* |
| --- | --- | --- | --- | --- | --- | --- |
| *Acidithiobacillus* | 2.18 | 1.000 |  |  |  |  |
| *Leptospirillum* | 2.30 | -0.144 (0.12) | 1.000 |  |  |  |
| *Ferrovum* | 2.98 | 0.020 (0.8) | 0.000 | 1.000 |  |  |
| *Ferritrophicum* | 3.89 | 0.000 | 0.000 | 0.102 (0.9) | 1.000 |  |
| *Gallionella* | 3.97 | 0.000 | 0.000 | 0.000 | -0.103 (0.1) | 1.000 |
| *Sideroxydans* | 4.50 | 0.012 (2.3) | 0.017 (2.2) | 0.000 | -0.068 (0.6) | -0.217 (0.5) |

**Table 5.**
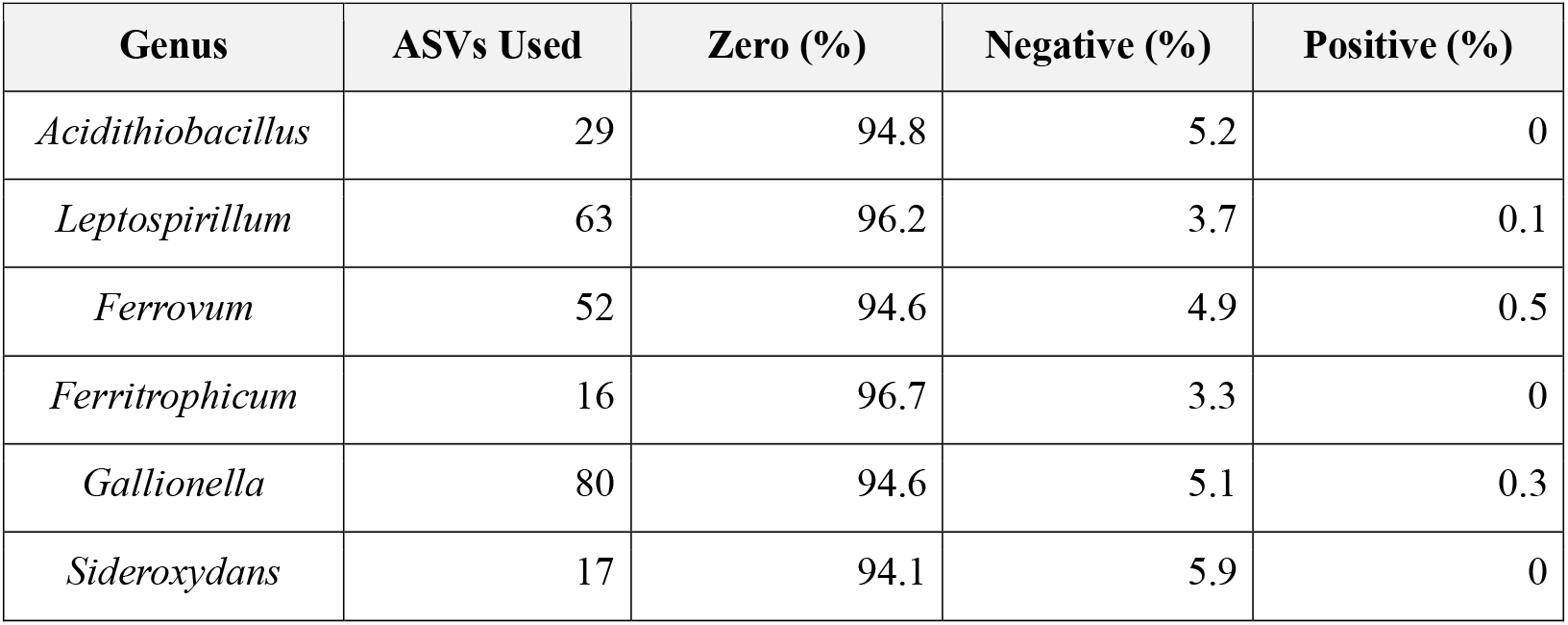
Summary of correlation coefficients for taxa within the same genus. Each column represents the percentage of ASVs within the genus that had no (zero), negative, or positive relationships with other members of the genus. Only taxa present in >3 samples were used.

## Discussion

AMD communities across the globe and spanning a wide variety of geochemical environments shared several key taxa. The core microbiome (those present in >75% of samples) contains Fe(II) oxidizing genera like *Ferrovum* [51], *Metallibacterium* which contains at least one Fe(II) reducing taxon [52] and *Acidibacter* [53], and the genus *Acidiferrobacter*, members of which are capable of iron and sulfur oxidation [54]. *Acidithiobacillus*, which typically cycles iron or other metals or sulfur is also a core taxon [55–58]. However, no ASVs of any genus were present in more than 34% of communities. Further only 46 ASVs were present in 20% or more of the communities. This suggests that while genera are commonly shared across AMD sites, individual species within common genera have narrower distributions.

### Distribution of Fe(II) oxidizing genera

Neutral models provide the opportunity to determine if the community composition is largely determined by stochastic processes like birth, death, immigration rates, or drift or deterministic processes like environmental filtering, competition, or by dispersal limitation. Many genera (44.3%) appear as frequently as expected (Figure 1A and Table 2), suggesting their distribution is controlled by stochastic or random processes like population drift rather than in response to environmental parameters. The majority (52.2%) of genera appear more frequently than predicted by the neutral model (Figure 1A and Table 2) consistent with positive selection, likely through environmental filtering. Both pH and high concentrations of metals may be filtering the taxa that can inhabit a specific AMD environment because they both impose physiological stress [e.g., 59, 60].

All of the major Fe(II) oxidizing genera (*Acidithiobacillus, Gallionella, Ferrovum, Leptospirillum, Ferrotrophicum*, and *Sideroxydans*) appeared less frequently than expected. Because these appear less frequently than expected, dispersal limitation or deterministic processes including environmental filtering or competitive interactions likely impact the distribution of Fe(II) oxidizing genera. Dispersal limitation is difficult to establish using 16S rRNA gene data because it does not provide the resolution necessary to identify biogeographic patterns. However, the “m” parameter, or immigration rate from the neutral model provides insight into dispersal into a community. In these models, when m = 1, every vacancy in the community created by death is filled by an immigrant. When m = 0, all vacancies are filled from the local community (i.e. by birth) [34]. In the > 300 AMD communities analyzed here, the m was 5.10^-5^. This low m values indicates AMD communities are dispersal limited: nearly every vacancy is filled from within the AMD community and there are few immigrants. Dispersal limitation can prevent more competitive species from establishing themselves in a population and makes the community composition more dependent on drift, a stochastic process [18], though the exact rate that causes stochasticity to dominate is unclear [18, 61, 62].

Both environmental filtering and ecological interactions like competition can cause microbial taxa to appear more or less frequently than predicted by neutral population models. Previous work has shown that the abundance of major Fe(II) oxidizing genera is controlled by environmental filtering, either by pH [5] or by a combination of pH and concentration of Fe(II) [19, 20]. In our dataset, pH is a strong environmental filter for acidophilic Fe(II) oxidizing taxa; it explains a significant portion of variation in Fe(II) oxidizing genera in the ∼300 samples we analyzed (Table 2). Consistent with previous studies, the genera *Leptospirillum* and *Acidithiobacillus* inhabit lower pH environments (pH optima 2.02 and 1.86 respectively) than *Ferrovum* which has a pH optimum of 3.12. *Ferritrophicum, Gallionella*, and *Sideroxydans* inhabit more moderate pH environments (pH optima of 3.89, 3.97 and 4.5 respectively). Because pH 4.5 is the highest pH considered in our dataset, the full range of *Sideroxydans* growth is likely outside the parameters of our dataset. In the laboratory *Sideroxydans* grows optimally at pH 6 [63]. Similarly, *Ferritrophicum* grew optimally at pH > 4.5 in the laboratory [43]; still, our data indicate they can be (relatively) abundant at lower pH.

While pH optima are evident in our data set, acidophilic Fe(II) oxidizing genera have considerable overlap in the pH in which they can persist (Figure 2). To test if these taxa are potential competitors, we used a graphical lasso on the residuals to determine if the genera covary. When the effect of pH was removed, the abundance of several Fe(II) oxidizing genera were negatively correlated indicative of a competitive interaction and the taxa that compete have similar pH optima. For example, *Leptospirillum* and *Acidithiobacillus* have pH optima near pH 2 and have a strong negative covariance. Similarly, *Ferritrophicum, Gallionella* and *Sideroxydans* are negatively correlated and appear to be competing for the Fe(II) oxidizing niche at pH 3.8 and above (Table 3, Figure 2). Taxa that do not have similar pH optima appear to either have zero correlation or in some cases, have a positive covariance (Table 4). Those that are positively correlated and thus unlikely to compete have pH optima that are separated by 0.8 pH units or more. This suggests that competition is limited to genera that are optimized for specific environmental conditions (e.g. pH).

### Distribution of Fe(II) oxidizing species

Individual Fe(II) oxidizing ASVs appear to be shaped by different forces than the genera as a whole. Although all Fe(II) oxidizing genera appeared less frequently than predicted by the neutral model, individual ASVs within each genus vary in whether they deviate from the neutral model (Table 2). This discrepancy between ASV and genus frequency indicates that some factors influence individual species that are not evident at the genus level and that intra-genus diversity may be important in AMD environments. The abundance of all genera had significant relationship with pH. In contrast, a smaller fraction (13.7%-47.0%) of ASVs had a significant relationship between ASV abundance and pH (for ASVs present in >3 sites; Table 3). Further, the ASVs that had a significant relationship between abundance and pH had a lower proportion, on average, of their variability explained by pH compared to the genus (Table 3) suggesting that pH plays a smaller role in structuring the abundance of the individual ASVs. Intra-genus interactions also do not appear to play an important role in the distribution of Fe(II) oxidizing ASVs. Approximately half of the ASVs are found at fewer than 3 sites and of those that do appear at >3 sites, >90% are not correlated (positively or negatively) with other taxa within the same genus (Table 5). Dispersal limitation appears to structure the distribution of Fe(II) oxidizing ASVs. The sparse distribution of ASVs and low m are both consistent with dispersal limitation at the ASV level. This pattern contrasts with the broad distribution at the genus level. Thus, across the range of pH observed in our study (pH < 5.0) dispersal limitation is the dominant factor in determining which Fe(II) oxidizing species are abundant. Because there is considerable variation in the metabolic potential of species within a single genus (e.g., with respect to nitrogen fixation, motility, and chemotaxis [22, 30]), this dispersal limitation may strongly influence how an ecosystem functions.

The observation that dispersal limitation plays an important role in the distribution of Fe(II) oxidizing species suggests seeding is a reasonable bioremediation strategy for AMD. On a first order level, target species should be selected based on the broad geochemical context – for example, only species within *Leptospirillum* or *Acidithiobacillus* should be selected for remediation of environments where the pH is <2.5. Then the optimal species could be selected based on their Fe(II) oxidation rate, or another desirable feature. Because dispersal is what is limiting their distribution, seeding efforts should allow the taxon to establish itself, though repeated seeding may be necessary. Once established, limited dispersal should help to maintain the new community composition, leading to a long-term increase in Fe(II) oxidation rate.

More broadly, variability in success in seeding efforts may result from variability in the factors that control the composition of microbial communities. Seeding communities that are primarily influenced by dispersal limitation is far more likely to succeed than those driven by environmental filtering or competitive interactions. Future seeding efforts should examine the ecological controllers of community composition in their chosen environment and use this information to inform their endeavors.

## Data Availability Statement

The datasets analyzed during the current study are available in the NCBI Sequence Read Archive under the accession numbers listed in Table 1.

## Acknowledgements

This work was supported by a grant from the National Science Foundation (2345568) awarded to Grettenberger and Hamilton. ChatGPT was used to aid the authors in writing and troubleshooting R code in sections where the statistical processes needed to be applied to all taxa and for generating figures in ggplot2.

## Works Cited

1. Druschel GK et al. Acid mine drainage biogeochemistry at Iron Mountain, California. Geochem T 2004;5:13. 10.1186/1467-4866-5-13

2. Johnson DB. Acidophilic microbial communities: Candidates for bioremediation of acidic mine effluents. Int Biodeter Biodegr 1995;35:41–58. 10.1016/0964-8305(95)00065-d

3. Grettenberger CL et al. Efficient Low-pH Iron Removal by a Microbial Iron Oxide Mound Ecosystem at Scalp Level Run. Appl Environ Microb 2017;83:e00015–17. 10.1128/aem.00015-17

4. Qiu G et al. Archaeal diversity in acid mine drainage from Dabaoshan Mine, China. J Basic Microb 2008;48:401–409. 10.1002/jobm.200800002

5. Kuang J-L et al. Contemporary environmental variation determines microbial diversity patterns in acid mine drainage. Isme J 2013;7:1038. 10.1038/ismej.2012.139

6. Johnson DB, Hallberg KB. Acid mine drainage remediation options: a review. Sci Total Environ 2005;338:3–14. 10.1016/j.scitotenv.2004.09.002

7. Hegler F et al. Physiology of phototrophic iron(II)_□oxidizing bacteria: implications for modern and ancient environments. FEMS Microbiol Ecol 2008;66:250–260. 10.1111/j.1574-6941.2008.00592.x

8. Johnson DB, Kanao T, Hedrich S. Redox Transformations of Iron at Extremely Low pH: Fundamental and Applied Aspects. Front Microbiol 2012;3:96. 10.3389/fmicb.2012.00096

9. Hvas CL et al. Fecal Microbiota Transplantation Is Superior to Fidaxomicin for Treatment of Recurrent Clostridium difficile Infection. Gastroenterology 2019;156:1324-1332.e3. 10.1053/j.gastro.2018.12.019

10. Mattila E et al. Fecal Transplantation, Through Colonoscopy, Is Effective Therapy for Recurrent Clostridium difficile Infection. Gastroenterology 2012;142:490–496. 10.1053/j.gastro.2011.11.037

11. Kassam Z et al. Fecal microbiota transplantation for Clostridium difficile infection: systematic review and meta-analysis. Am J Gastroenterol 2013;108:500–8. 10.1038/ajg.2013.59

12. Leahy JG, Colwell RR. Microbial degradation of hydrocarbons in the environment. Microbiol Rev 1990;54:305–15. 10.1128/mr.54.3.305-315.1990

13. Arif I, Batool M, Schenk PM. Plant Microbiome Engineering: Expected Benefits for Improved Crop Growth and Resilience. Trends Biotechnol 2020;38:1385–1396. 10.1016/j.tibtech.2020.04.015

14. Peng X, Bruns MA. Cyanobacterial Soil Surface Consortia Mediate N Cycle Processes in Agroecosystems. Front Environ Sci 2019;6:156. 10.3389/fenvs.2018.00156

15. Loakasikarn T et al. Effect of seeding materials on organic matter degradation and microbial community succession during model organic waste composting. Biocatal Agric Biotechnology 2021;37:102182. 10.1016/j.bcab.2021.102182

16. Friedman J, Higgins LM, Gore J. Community structure follows simple assembly rules in microbial microcosms. Nat Ecol Evol 2017;1:0109. 10.1038/s41559-017-0109

17. Goldford JE et al. Emergent simplicity in microbial community assembly. Science 2018;361:469–474. 10.1126/science.aat1168

18. Evans S, Martiny JBH, Allison SD. Effects of dispersal and selection on stochastic assembly in microbial communities. ISME J 2017;11:176–185. 10.1038/ismej.2016.96

19. Macalady JL et al. Energy, ecology and the distribution of microbial life. Philos Trans R Soc B Biol Sci 2013;368:20120383. 10.1098/rstb.2012.0383

20. Jones DS et al. Geochemical Niches of Iron-Oxidizing Acidophiles in Acidic Coal Mine Drainage. Appl Environ Microb 2015;81:1242–1250. 10.1128/aem.02919-14

21. Havig JR, Grettenberger C, Hamilton TL. Geochemistry and microbial community composition across a range of acid mine drainage impact and implications for the Neoarchean_□Paleoproterozoic transition. J Geophys Res Biogeosciences 2017;122:1404–1422. 10.1002/2016jg003594

22. Grettenberger CL, Havig JR, Hamilton TL. Metabolic diversity and co-occurrence of multiple Ferrovum species at an acid mine drainage site. Bmc Microbiol 2020;20:119. 10.1186/s12866-020-01768-w

23. Apprill A et al. Minor revision to V4 region SSU rRNA 806R gene primer greatly increases detection of SAR11 bacterioplankton. Aquat Microb Ecol 2015;75:129–137. 10.3354/ame01753

24. Parada AE, Needham DM, Fuhrman JA. Every base matters: assessing small subunit rRNA primers for marine microbiomes with mock communities, time series and global field samples. Environ Microbiol 2016;18:1403–1414. 10.1111/1462-2920.13023

25. She Z et al. Decadal evolution of an acidic pit lake: Insights into the biogeochemical impacts of microbial community succession. Water Res 2023;243:120415. 10.1016/j.watres.2023.120415

26. Callahan BJ et al. DADA2: High-resolution sample inference from Illumina amplicon data. Nat Methods 2016;13:nmeth.3869. 10.1038/nmeth.3869

27. Quast C et al. The SILVA ribosomal RNA gene database project: improved data processing and web-based tools. Nucleic Acids Res 2013;41:D590–D596. 10.1093/nar/gks1219

28. McMurdie PJ, Holmes S. phyloseq: An R Package for Reproducible Interactive Analysis and Graphics of Microbiome Census Data. Plos One 2013;8:e61217. 10.1371/journal.pone.0061217

29. Marcon E, Hérault B. entropart□: An R Package to Measure and Partition Diversity. J Stat Softw 2015;67. 10.18637/jss.v067.i08

30. Grettenberger CL, Macalady JL, Hamilton TL. Metabolic diversity of the Ferrovales and potential contributions to iron oxidation. bioRxiv 2025;2025.11.03.686282. 10.1101/2025.11.03.686282

31. Team Rs. RStudio: Integrated Development for R. Boston, MA: RStudio, Inc., 2019.

32. Team” “R Core. R: A Language and Environment for Statistical Computing. R Foundation for Statistical Computing, 2023.

33. Russell88. Russel88/MicEco: Various functions for microbial community data. 2022.

34. Sloan WT et al. Quantifying the roles of immigration and chance in shaping prokaryote community structure. Environ Microbiol 2006;8:732–740. 10.1111/j.1462-2920.2005.00956.x

35. Burns AR et al. Contribution of neutral processes to the assembly of gut microbial communities in the zebrafish over host development. ISME J 2015;10:655–664. 10.1038/ismej.2015.142

36. Brown JF et al. Application of a depositional facies model to an acid mine drainage site. Appl Environ Microb 2010;77:545–54. 10.1128/aem.01550-10

37. Sheng Y et al. Bioreactors for low-pH iron( ii ) oxidation remove considerable amounts of total iron. Rsc Adv 2017;7:35962–35972. 10.1039/c7ra03717a

38. Senko JM et al. Characterization of Fe(II) oxidizing bacterial activities and communities at two acidic Appalachian coalmine drainage-impacted sites. Isme J 2008;2:1134. 10.1038/ismej.2008.60

39. Auld RR et al. Characterization of the microbial acid mine drainage microbial community using culturing and direct sequencing techniques. J Microbiol Methods 2013;93:108–115. 10.1016/j.mimet.2013.01.023

40. Sheng Y et al. Geochemical and Temporal Influences on the Enrichment of Acidophilic Iron-Oxidizing Bacterial Communities. Appl Environ Microb 2016;82:3611–3621. 10.1128/aem.00917-16

41. Hallberg KB et al. Macroscopic Streamer Growths in Acidic, Metal-Rich Mine Waters in North Wales Consist of Novel and Remarkably Simple Bacterial Communities. Appl Environ Microb 2006;72:2022–2030. 10.1128/aem.72.3.2022-2030.2006

42. Ramos-Perez D et al. Changes in the prokaryotic diversity in response to hydrochemical variations during an acid mine drainage passive treatment. Sci Total Environ 2022;842:156629. 10.1016/j.scitotenv.2022.156629

43. Weiss JV et al. Characterization of Neutrophilic Fe(II)-Oxidizing Bacteria Isolated from the Rhizosphere of Wetland Plants and Description of Ferritrophicum radicicola gen. nov. sp. nov., and Sideroxydans paludicola sp. nov. Geomicrobiol J 2007;24:559–570. 10.1080/01490450701670152

44. Emerson D et al. Comparative genomics of freshwater Fe-oxidizing bacteria: implications for physiology, ecology, and systematics. Front Microbiol 2013;4:254. 10.3389/fmicb.2013.00254

45. Bredberg K, Karlsson HT, Holst O. Reduction of vanadium(V) with Acidithiobacillus ferrooxidans and Acidithiobacillus thiooxidans. Bioresour Technol 2003;92:93–6. 10.1016/j.biortech.2003.08.004

46. Yin H et al. Whole-genome sequencing reveals novel insights into sulfur oxidation in the extremophile Acidithiobacillus thiooxidans. BMC Microbiol 2014;14:179–179. 10.1186/1471-2180-14-179

47. Boogaart KG van den, Tolosana-Delgado R. “compositions”: A unified R package to analyze compositional data. Comput Geosci 2008;34:320–338. 10.1016/j.cageo.2006.11.017

48. Wood S. mgcv: GAMs and generalized ridge regression for R. R News 2001;1:20–25.

49. Friedman J, Hastie T, Tibshirani R. Sparse inverse covariance estimation with the graphical lasso. Biostatistics 2007;9:432–441. 10.1093/biostatistics/kxm045

50. Zhao T et al. The huge Package for High-dimensional Undirected Graph Estimation in R. J Mach Learn Res_□: JMLR 2012;13:1059–1062.

51. Johnson DB, Hallberg KB, Hedrich S. Uncovering a Microbial Enigma: Isolation and Characterization of the Streamer-Generating, Iron-Oxidizing, Acidophilic Bacterium “Ferrovum myxofaciens.” Appl Environ Microb 2014;80:672–680. 10.1128/aem.03230-13

52. Ziegler S et al. Metallibacterium scheffleri gen. nov., sp. nov., an alkalinizing gammaproteobacterium isolated from an acidic biofilm. Int J Syst Evol Microbiol 2013;63:1499– 1504. 10.1099/ijs.0.042986-0

53. Falagán C, Johnson DB. Acidibacter ferrireducens gen. nov., sp. nov.: an acidophilic ferric iron-reducing gammaproteobacterium. Extremophiles 2014;18:1067–1073. 10.1007/s00792-014-0684-3

54. Hallberg KB, Hedrich S, Johnson DB. Acidiferrobacter thiooxydans, gen. nov. sp. nov.; an acidophilic, thermo-tolerant, facultatively anaerobic iron- and sulfur-oxidizer of the family Ectothiorhodospiraceae. Extremophiles 2011;15:271–279. 10.1007/s00792-011-0359-2

55. Castelle C et al. A New Iron-oxidizing/O2-reducing Supercomplex Spanning Both Inner and Outer Membranes, Isolated from the Extreme Acidophile Acidithiobacillus ferrooxidans. J Biol Chem 2008;283:25803–25811. 10.1074/jbc.m802496200

56. Hallberg KB, González-Toril E, Johnson DB. Acidithiobacillus ferrivorans, sp. nov.; facultatively anaerobic, psychrotolerant iron-, and sulfur-oxidizing acidophiles isolated from metal mine-impacted environments. Extremophiles 2010;14:9–19. 10.1007/s00792-009-0282-y

57. Quatrini R et al. Extending the models for iron and sulfur oxidation in the extreme Acidophile Acidithiobacillus ferrooxidans. BMC Genom 2009;10:394–394. 10.1186/1471-2164-10-394

58. Mangold S et al. Sulfur Metabolism in the Extreme Acidophile Acidithiobacillus Caldus. Front Microbiol 2011;2:17. 10.3389/fmicb.2011.00017

59. Guan N, Liu L. Microbial response to acid stress: mechanisms and applications. Appl Microbiol Biotechnol 2020;104:51–65. 10.1007/s00253-019-10226-1

60. Prabhakaran P, Ashraf MA, Aqma WS. Microbial stress response to heavy metals in the environment. RSC Adv 2016;6:109862–109877. 10.1039/c6ra10966g

61. Declerck SAJ et al. Effects of patch connectivity and heterogeneity on metacommunity structure of planktonic bacteria and viruses. ISME J 2012;7:533–542. 10.1038/ismej.2012.138

62. Lindström ES, Östman Ö. The Importance of Dispersal for Bacterial Community Composition and Functioning. PLoS ONE 2011;6:e25883. 10.1371/journal.pone.0025883

63. Kato S et al. Sideroxyarcus emersonii gen. nov. sp. nov., a neutrophilic, microaerobic iron- and thiosulfate-oxidizing bacterium isolated from iron-rich wetland sediment. Int J Syst Evol Microbiol 2022;72. 10.1099/ijsem.0.005347

64. Burkartová K et al. Distinct microbial communities supported by iron oxidation. Environ Microbiol 2024;26:e16706. 10.1111/1462-2920.16706

65. Grettenberger CL, Hamilton TL. Metagenome-Assembled Genomes of Novel Taxa from an Acid Mine Drainage Environment. Appl Environ Microb 2021;87:e0077221. 10.1128/aem.00772-21

66. Hao Y-Q et al. Microbial biogeography of acid mine drainage sediments at a regional scale across southern China. FEMS Microbiol Ecol 2022;98:fiac002. 10.1093/femsec/fiac002

67. Abramov SM et al. Biogeochemical Niches of Fe-Cycling Communities Influencing Heavy Metal Transport along the Rio Tinto, Spain. Appl Environ Microbiol 2021;88:e02290–21. 10.1128/aem.02290-21

68. She Z et al. Vertical environmental gradient drives prokaryotic microbial community assembly and species coexistence in a stratified acid mine drainage lake. Water Res 2021;206:117739. 10.1016/j.watres.2021.117739

69. Wang M et al. Strong succession in prokaryotic association networks and community assembly mechanisms in an acid mine drainage-impacted riverine ecosystem. Water Res 2023;243:120343. 10.1016/j.watres.2023.120343

70. Habe H et al. Distinctive microbial communities linked with pH and heavy metals in mine drainages across all regions of Japan. Sci Total Environ 2025;975:179222. 10.1016/j.scitotenv.2025.179222

71. Ayala-Muñoz D et al. Metagenomic and Metatranscriptomic Study of Microbial Metal Resistance in an Acidic Pit Lake. Microorg 2020;8:1350. 10.3390/microorganisms8091350

