## Supplemental Table 1 for "Environmental filtering, dispersal limitation, and competition control the distribution of acidophilic iron oxidizers"

**Supplemental Table 1.** Percentage of variance in the abundance (R^2^) of members of the genus *Acidithiobacillus* that is explained by pH using a generalized additive model, the significance of the relationship (p-value) and the estimated degrees of freedom (EDF) for the chosen model. The pH optimum for the species and the average relative abundance are indicated in the final two columns.

| **Species** | **R^2^** | **p-value** | **EDF** | **pH optimum** | **Average Relative Abundance (%)** |
| --- | --- | --- | --- | --- | --- |
| *A. ferrooxidans* | 0.05 | <0.01 | 2.2 | 2.19 | 4.0 |
| Unclassified *Acidithiobacillus* | 0.04 | <0.01 | 1.3 | 2.19 | 0.5 |
| *A. caldus* | 0.01 | 0.02 | 1.0 | 2.19 | 0.8 |
| *A. ferrovorans* | 0.02 | 0.07 | 1.9 | 2.96 | 0.72 |
| *A. ferriphilus* | 0.08 | <0.01 | 2.4 | 3.85 | <0.01 |
| *A. albertensis* | 0.10 | <0.01 | 2.4 | 3.99 | <0.01 |
| *A. ferridurans* | 0.08 | <0.01 | 2.9 | 4.05 | <0.01 |
| *A. thiooxidans* | 0.10 | <0.01 | 1.7 | 4.50 | 0.03 |
